# A Cell-Based Assay to Categorize Variants of Human Succinate Semialdehyde Dehydrogenase Associated with Autism Spectrum Disorder

**DOI:** 10.64898/2026.01.15.699751

**Authors:** Barry P Young, Isaac Robinson, Jessica Schmitt, Brian Wu, Jie Liu, Jesse T Chao, Sanja Rogic, Paul Pavlidis, Douglas W. Allan, Christopher J R Loewen

## Abstract

Many genes are implicated in autism spectrum disorder (ASD) as a result of sequencing the genomes of individuals with ASD, however in most cases it remains unclear which genes are playing causative roles. *ALDH5A1*, which encodes the enzyme succinate semialdehyde dehydrogenase (SSADH), an important regulator of GABA metabolism, is causative in the syndrome SSADH deficiency and is also implicated in ASD. However, it is unknown how variants found in ASD affect SSADH function. We developed a yeast growth assay that models SSADH deficiency to quantify the functional impact of seven *ALDH5A1* variants found in ASD. In this assay, expression of human *ALDH5A1* partially complemented the growth defect caused by deletion of the *ALDH5A1* ortholog *UGA2*. Using growth rate measurements, we calculated functional scores for 27 variants divided into calibration and ASD test variant groups. Functional scores for benign and pathogenic calibration variants segregated accordingly, validating the assay, while ASD variants displayed a range of activities from complete loss to normal function. Comparisons with published enzymatic assays and computational predictions showed broad agreement, while also identifying some limitations of these approaches.

**SUMMARY STATEMENT:** We introduce a simple yeast assay to measure how autism-linked gene variants in *ALDH5A1*, which encodes the enzyme succinate semialdehyde dehydrogenase, affect its function, helping clarifying its role in autism.

## INTRODUCTION

Autism spectrum disorder (ASD) is a common neurodevelopmental condition characterized by persistent difficulties in social communication and interaction, along with restricted, repetitive behaviours and interests. Globally, ASD affects about 1 in 100 children, with similar estimates reported in multiple epidemiological surveys (Zeidan et al., 2022). Genetic factors contribute substantially to ASD risk, with heritability estimates between 70% and 90% in twin studies, and hundreds of genes now implicated through rare and common variants (Genovese and Butler, 2023). The genetic architecture spans *de novo* variants, rare inherited variants and common variants, reflecting the complexity and heterogeneity of ASD.

Syndromic ASD is defined when ASD is part of a broader medical condition, often associated with variants in a single gene. One gene which has been associated with syndromic ASD is *ALDH5A1*, which encodes succinate semialdehyde dehydrogenase (SSADH). Pathogenic variants in *ALDH5A1* result in SSADH deficiency, a rare inherited metabolic autosomal disorder, caused by bi-allelic mutations. Estimates of worldwide prevalence vary between 1 in 223,000 to 1 in 564,000 (Glinton et al., 2024). Clinically, the syndrome presents as a range of symptoms including aggression, ataxia, hyperkinetic behaviour, hypotonia, intellectual disability, moderate-to-severe developmental delays, psychiatric disorders and seizures.

Individuals affected by SSADH deficiency exhibit impaired degradation of the inhibitory neurotransmitter gamma-aminobutyric acid (GABA). The first stage of GABA catabolism involves the action of GABA transaminase, which catalyzes the conversion of GABA to succinate semialdehyde (SSA) (Figure 1A). In unaffected individuals, succinate semialdehyde is then converted to succinate by SSADH, with the concomitant conversion of NAD(P)^+^ to NAD(P)H^+^. However, in SSADH deficiency patients, this step is impaired and results in accumulation of metabolites including GABA, γ-guanidinobutyrate and gammahydroxybutyrate (GHB); the latter of which can be detected at elevated levels in urine, plasma and cerebrospinal fluid and is used as a diagnostic tool in SSADH deficiency.

**Figure 1:**
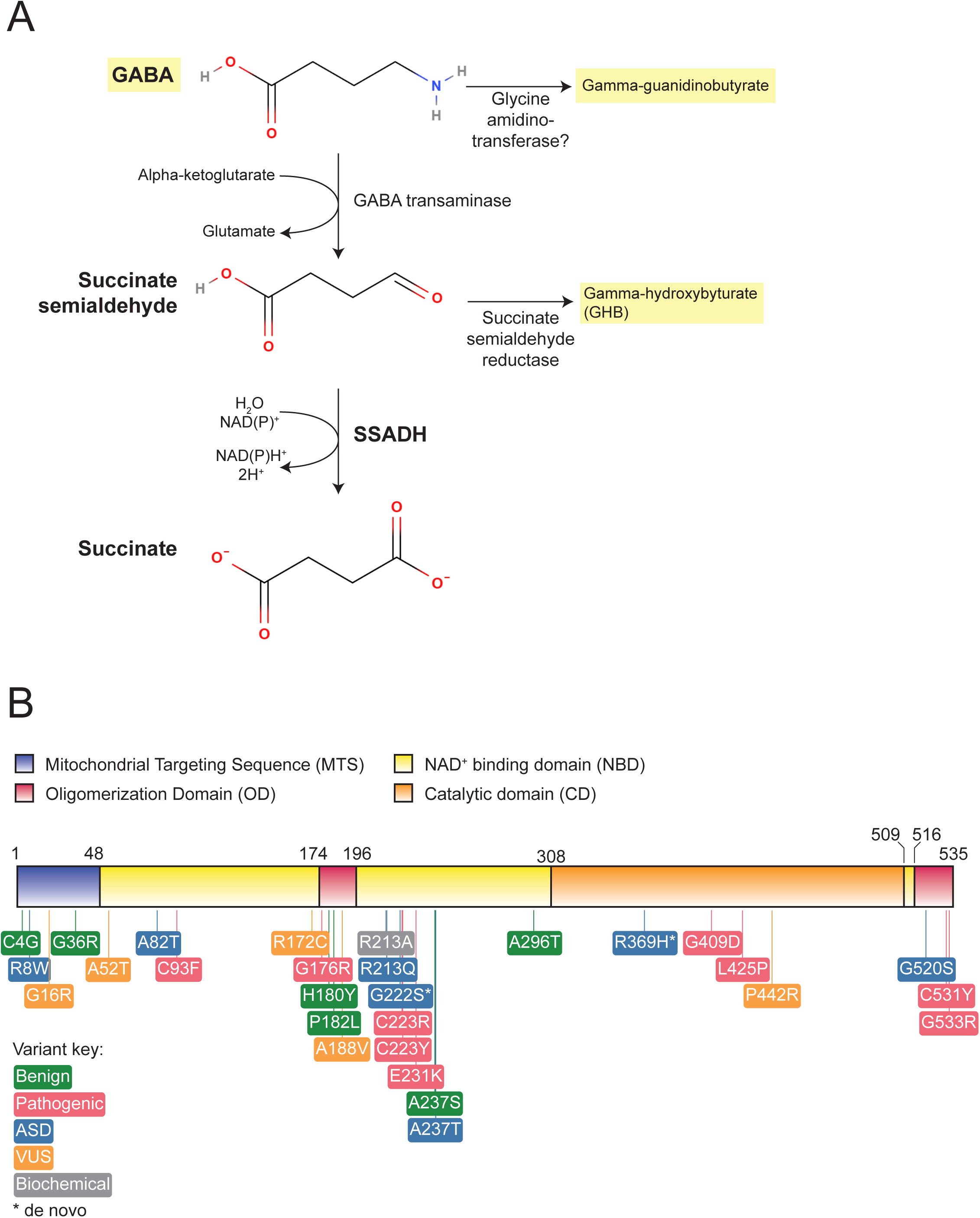
Function and structure of SSADH. (A) SSADH catalyzes the conversion of succinate semialdehyde to succinate with the concomitant production of NAD(P)H from NAD(P)^+^. In the absence of SSADH function, GABA, gamma-hydroxybutyrate (GHB) and gamma-guanidinobutryate (highlighted in yellow) are found at elevated levels. (B) Domain structure of human SSADH showing the position of the variants examined in this study.

The cerebral accumulation of GABA and GHB disrupts the balance between GABAergic (excitatory) and glutamatergic (inhibitory) neurotransmission. ASD has been associated with impairment of the GABAergic nervous system, and ASD is prevalent in SSADH deficiency (Tokatly Latzer et al., 2023). Interestingly, SSADH deficiency with ASD is often associated with lower plasma levels of GABA and GHB than SSADH deficiency without ASD (Tokatly Latzer et al., 2023). While there are clinical means to diagnose SSADH deficiency, there are limited predictive tools available to assess *ALDH5A1* variants. Traditionally, the enzymatic activity of SSADH variants has been quantified using an *in vitro* assay based on the change in NAD^+^ fluorescence upon addition of SSA to extracts made from clinical samples or transfected cells expressing the variant of interest (Akaboshi et al., 2003; Blasi et al., 2002; Gibson et al., 1991; Pop et al., 2020).

The budding yeast *Saccharomyces cerevisiae* has been successfully used for high-throughput analysis of a number of human disease genes and their variants by our lab and others (Chen et al., 2025; Kachroo et al., 2022; Mighell et al., 2018; Post et al., 2020; Sun et al., 2016; Young et al., 2020). Here we present an alternative *in vivo* assay based on the ability of human *ALDH5A1* to complement the activity of the orthologous yeast *UGA2* gene. Using this assay, we were able to accurately measure the function of a large number of variants with high precision and reproducibility, including seven variants found in ASD individuals. By comparison to known benign and pathogenic calibration variants we were able to classify all seven as either benign or pathogenic. Most of these were variants of uncertain significance (VUS) without a clear clinical interpretation. This approach demonstrates a rapid, high-throughput method of evaluating *ALDH5A1* variants that will serve to complement existing genetic and clinical testing for SSADH deficiency and ASD.

## METHODS

### Variant aggregation, annotation, and prioritization

We utilized our in-house computational pipeline to aggregate *ALDH5A1* missense variants from ClinVar and VariCarta and perform comprehensive annotation. We collected ClinVar variants (downloaded in January 2022) that have at least one gold review star, i.e., the review status (CLNREVSTAT) has to have one of the following designations: “criteria provided, single submitter,” “criteria provided, conflicting interpretations,” “criteria provided, multiple submitters, no conflicts” or “reviewed by expert panel.” For each of these variants, we also obtain its interpretation (clinical significance; CLNSIG) and reported condition (disease name; CLNDN). The ASD missense variants were obtained from VariCarta, a database of variants found in individuals diagnosed with ASD and reported in the scientific literature, that we created (Belmadani et al., 2019). We annotated the aggregated variants using the Ensembl VEP tool (McLaren et al., 2016). We also added protein domain annotation.

### Plasmids

A plasmid containing the coding sequence of human *ALDH5A1* (ENSMBL sequence ENST00000357578) in a Gateway entry vector, pTwist-ENTR-*ALDH5A1*, was codon-optimized for expression in yeast and ordered from Twist Biosciences. For expression in yeast, *ALDH5A1* and variants were transferred to the Gateway destination vector pAG416-GPD-ccdB (low copy, Addgene plasmid # 14148) by Gateway cloning (ThermoFisher). The plasmid was a gift from Susan Lindquist (Alberti et al., 2007).

### Site-directed mutagenesis

Primers for mutagenesis were designed using the NEBaseChanger web site (https://nebasechanger.neb.com/). Mutagenesis reactions were performed using the Q5 PCR kit according to manufacturer’s instructions in 25 µl reactions with 0.1 ng of pTwist-ENTR-*ALDH5A1* as template. Following confirmation of amplification, 1 µl of PCR products were treated with 1 µl DpnI, 1 µl T4 DNA Ligase, 1 µl polynucleotide kinase in T4 ligase buffer (all reagents from NEB) for one hour at room temperature. The reaction was transformed into *E. coli* and mutant plasmids were confirmed by whole plasmid sequencing (Plasmidsaurus Inc). For expression in yeast, plasmids were transferred to pAG416-GPD-ccdB using the Gateway LR Clonase II kit (Thermo Fisher Scientific) according to manufacturer instructions. Correct recombination was verified by whole plasmid sequencing.

### Growth Media

GABA Assay Medium (GAM) contained 0.67% yeast nitrogen base (YNB) without ammonium sulphate, 2% dextrose, 0.1% monosodium glutamate (MSG), 0.1% gamma-aminobutyric acid (GABA), 0.02% histidine, 0.1% leucine and 0.02% methionine.

### Yeast liquid growth assays

Transformants were robotically arrayed on to Rich-uracil agar plates using a Singer RoToR HDA robot prior to each experimental run and incubated overnight at 30 °C. Cells were then robotically transferred to 96-well dishes with 200 µl of GAM media in each well. Plates were shaken at 750 rpm overnight at 30 °C, after which each 40 µl of each culture was added to 160 µl of fresh GAM media. Growth was monitored using a LogPhase-600 plate reader (Agilent) by measuring absorbance at 600 nm for 24 hours. The “Vmax” value reported by the plate reading software was used as a measure of maximal cell growth in subsequent analyses. All variants were measured together alongside controls in four separate experiments with twelve biological replicates of each plasmid.

### Statistical analysis of variant effects

Yeast growth constants (k) were analyzed using methods similar to Chen *et al*. and Post *et al*. (Chen et al., 2025; Post et al., 2020). Briefly, linear mixed-effects models were fit to each dataset using the lmerTest R package (Kuznetsova et al., 2017) to estimate the phenotypic effects of each variant compared to the *ALDH5A1*-Ref construct, treating experimental batches as random effects (for R scripts see Supplementary Methods 1). For visualization, values were standardized to the range [0–1] where 0 represents the mean phenotype measured for an empty vector, and 1 represents the mean phenotype with the *ALDH5A1*-Ref construct.

## RESULTS

### Selection of Variants

The purpose of this study was to develop a yeast assay to precisely measure the functional impact of ASD-associated variants in *ALDH5A1*. We first collected all known missense variants and used this as a source for variant selection (Table S1). In order to determine that our assay would provide clinically relevant functional information, we selected a number of variants with clear benign and pathogenic clinical classifications, which we refer to as calibration variants (Figure 1). In general, we selected calibration variants with the highest confidence in their annotation, specifically those with multiple submitters and no conflicts in interpretation in the ClinVar database (Landrum et al., 2018). Where possible, we aimed to include calibration variants across the four domains of the SSADH protein. Benign calibration variants, when assayed, should result in a range of values that correspond to the fully functional protein, while pathogenic calibration variants should indicate values associated with reduced function. Additionally we included a variant, R213A, which has been used in biochemical studies and results in a seven-fold reduction in SSADH activity as measured *in vitro* (Kim et al., 2009). For ASD variants we identified eight variants that were present identified in individuals with ASD and present in the VariCarta database (Belmadani et al., 2019) (Table 1). We were unable to make one variant, R25H, so in total we analyzed seven ASD variants. These seven variants were of unknown clinical significance with some also present in ClinVar as “uncertain significance” or “conflicting interpretation”. Finally, we included 5 additional variants (designated VUS) that were not found in individuals with ASD, but that were present ClinVar as “uncertain significance” or “conflicting interpretation”. In total, we generated 27 missense variants corresponding to four distinct classes; benign, pathogenic, ASD and VUS (Figure 1).

**Table 1.** List of ALDH5A1 ASD variants.

| Variant | Domain | References/Source |
| --- | --- | --- |
| R8W | MTS | (Leblond et al., 2019) |
| R25H | MTS | (Zhou et al., 2022) |
| A82T | NBD | (Yalcintepe et al., 2021) |
| R213Q | NBD | (Guo et al., 2018) |
| G222S | NBD | (Dou et al., 2017; Iossifov et al., 2014; Ji et al., 2016; Kosmicki et al., 2017; Krupp et al., 2017; Lim et al., 2017; Zhou et al., 2022) |
| A237T | NBD | (Stessman et al., 2017) |
| R369H | CD | (Fu et al., 2022; Kosmicki et al., 2017; Lim et al., 2017; Satterstrom et al., 2020; The DDD Study et al., 2014; Trost et al., 2022; Zhou et al., 2022) |
| G520S | NBD | (Stessman et al., 2017) |

### Human ALDH5A1 partially complements uga2Δ yeast cells

Given that the enzymatic function of SSADH is precisely defined, we reasoned that the human protein may be able to compensate for the absence of the yeast homologue of SSADH, Uga2. Previous studies have shown that *uga2Δ* yeast cells are unable to grow with GABA as their sole nitrogen source (Ramos et al., 1985). We therefore tested whether expression of *ALDH5A1* in *uga2Δ* cells could restore their ability to utilize GABA. We observed only a minor rescue of the *uga2Δ* phenotype, resulting in growth that was too slow to measure accurately in liquid growth assays (Figure 2A). However, we found that when *uga2Δ* cells were grown in the presence of both 0.1% glutamate and 0.1% GABA, a fitness advantage was observed in cells expressing *ALDH5A1*. We found that increasing the concentration of GABA further reduced the growth rate of *uga2Δ* cells, which was alleviated by expression of *ALDH5A1* (Figure 2B). This suggested that *ALDH5A1* was functioning by repairing the GABA degradation pathway leading to increased fitness.

**Figure 2:**
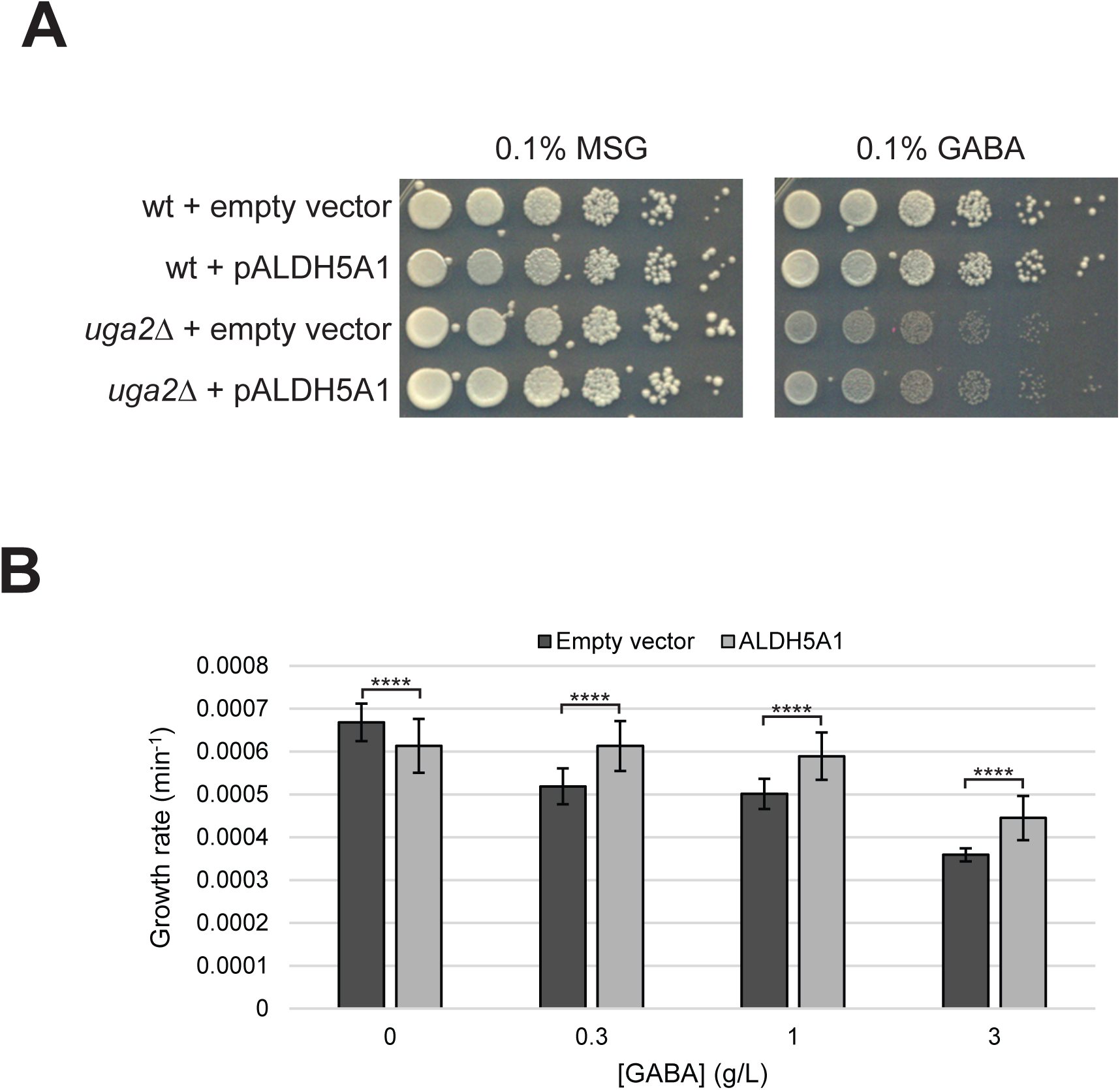
Complementation of *uga2Δ* yeast by human SSADH. (A) Growth of wild type and *uga2Δ* cells grown with glutamate (MSG) or GABA as nitrogen source. Cells were transformed with a plasmid expressing *ALDH5A1* or the corresponding empty vector. (B) Increasing GABA concentration in the presence of 0.1% glutamate causes a growth defect which can be alleviated by expression of *ALDH5A1*. Each bar represents six technical measurements of eight independent transformants. Error bars denote SD, **** denotes p<10^-7^.

### Functional assessment of ALDH5A1 variants

To further demonstrate that the observed phenotype in *uga2Δ* cells was directly due to expression of *ALDH5A1*, we measured the growth of this strain in GABA media while expressing a number of pathogenic and benign calibration variants of *ALDH5A1*. We reasoned that benign (*i.e.*, functional) variants should be better able to increase growth of *uga2Δ* cells than pathogenic (non-functional) variants. To precisely test this, we constructed four 96-well format arrays consisting of *uga2Δ* yeast transformed with a range of expression plasmids. We included the reference *ALDH5A1* sequence (*ALDH5A1*-Ref), the corresponding empty vector, the nine pathogenic variants, the six benign variants and the R2123A biochemical variant. In this array, we also included the seven ASD variants and the five VUS. Each plasmid was tested from twelve independent transformants. The entire experiment was repeated on four separate occasions. We measured the maximum rate of log-phase growth of each well in defined media containing 0.1% glutamate and 0.1% GABA.

The growth rates from these assays (Table S2) were then analyzed using a linear mixed-effects model (see Methods) to derive a scaled estimate of *ALDH5A1* function, with a value of 1 representing reference *ALDH5A1* function and a value of 0 indicating no function (Figure 3). We observed a clear separation between benign and pathogenic calibration variants. Of the pathogenic variants, eight out of nine scored below ∼0.61, while all benign variants scored above ∼0.77. Only pathogenic variants showed a significant decrease compared to *ALDH5A1*-Ref, whereas all benign variants were not significantly different. A single pathogenic variant in the OD domain (C531Y) appeared functional in this assay with a score of 0.83. We found that the ASD variants exhibited a range of activities ranging from complete loss of function to normal function. Of the 5 VUS, only P442R showed reduced function. Finally, we found that the biochemical variant R213A, which shows a seven-fold reduction in catalytic activity *in vitro* (Kim et al., 2009), had substantially reduced function, supporting that the assay primarily reported on SSADH enzymatic activity.

**Figure 3:**
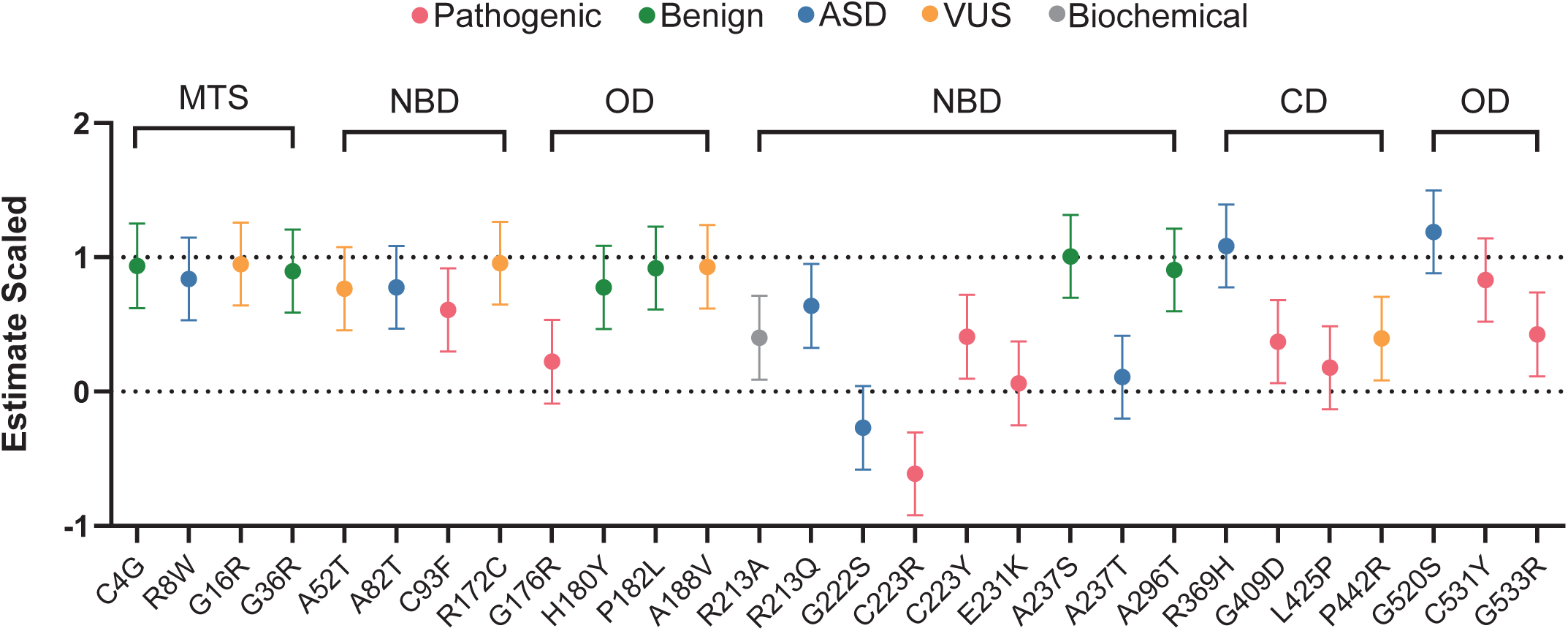
Estimated functional effects of *ALDH5A1* variants. Variants were expressed in *uga2Δ* yeast grown in 0.1% glutamate, 0.1% GABA and growth rates were analyzed using a linear mixed effects model. Data is normalized to reference *ALDH5A1* (=1) and empty vector (=0). Error bars indicate 95% confidence intervals.

### Comparison to published SSADH activity measurements

Previously, SSADH function has been measured using a fluorescent assay based on NAD^+^ consumption (See Table 2). Although we found broad overall agreement between this study and previously calculated values there were some notable differences. In particular, the pathogenic variants C93F, C223Y, G409D, C531Y and C533R appeared to be more functional in our yeast assay than *in vitro* assays. Conversely, the benign variants P182L and A237S appear to show substantial reductions in activity in *in vitro* assays whereas our assay indicated similar function as *ALDH5A1*-Ref.

**Table 2:** Comparison to in vitro measurements of SSADH activity.

| Variant | ClinVar Classification | This study | Others | Reference |
| --- | --- | --- | --- | --- |
| G36R | Benign | 0.897 | 0.87 | (Blasi et al., 2002) |
| C93F | Pathogenic | 0.609 | 0.03 | (Akaboshi et al., 2003) |
| G176R | Pathogenic | 0.223 | 0.01 | (Akaboshi et al., 2003) |
| H180Y | Benign | 0.777 | 0.825 | (Blasi et al., 2002) |
| P182L | Benign | 0.920 | 0.48 | (Blasi et al., 2002) |
| C223R | Pathogenic | -0.612 | 0.05 | (Pop et al., 2020) |
| C223Y | Pathogenic | 0.407 | 0.05 | (Akaboshi et al., 2003) |
| E231K | Pathogenic | 0.06 | 0.48, 0.42 | (Pop et al., 2020) |
| A237S | Benign | 1.007 | 0.651 | (Blasi et al., 2002) |
| A237T | VUS | 0.657 | 0.4, 0.36 | (Blasi et al., 2002) |
| G409D | Pathogenic | 0.372 | 0.01 | (Akaboshi et al., 2003) |
| C531Y | Pathogenic | 0.830 | 0.01 | (Pop et al., 2020) |
| C533R | Pathogenic | 0.427 | 0.01 | (Akaboshi et al., 2003) |

### Comparison to computational predictions

Computational methods offer an alternative path to classifying variants of unknown significance. While these techniques require significantly fewer resources, they can be limited in their predictive power (Molotkov et al., 2024). To compare our results to computational predictors, we first needed to translate our functional assay scores into predictions of pathogenicity. To do this we used the functional estimates for the variants from the model and the associated 95% confidence intervals (Figure 3). Variants whose 95% confidence intervals overlapped 1 indicating they were not significantly less functional than *ALDH5A1*-Ref were classified as likely benign, whereas variants whose 95% confidence intervals did not overlap 1 were classified as likely pathogenic (Figure 4). Out of the seven ASD variants, we classified three as likely pathogenic (R213Q, G222S and A237T) and four as likely benign (R8W, A82T, R369H and G520S). Of the five VUS, we classified only one as likely pathogenic (P442R) while the remaining four were classified as likely benign (G16R, A52T, R172C and A188V).

**Figure 4:**
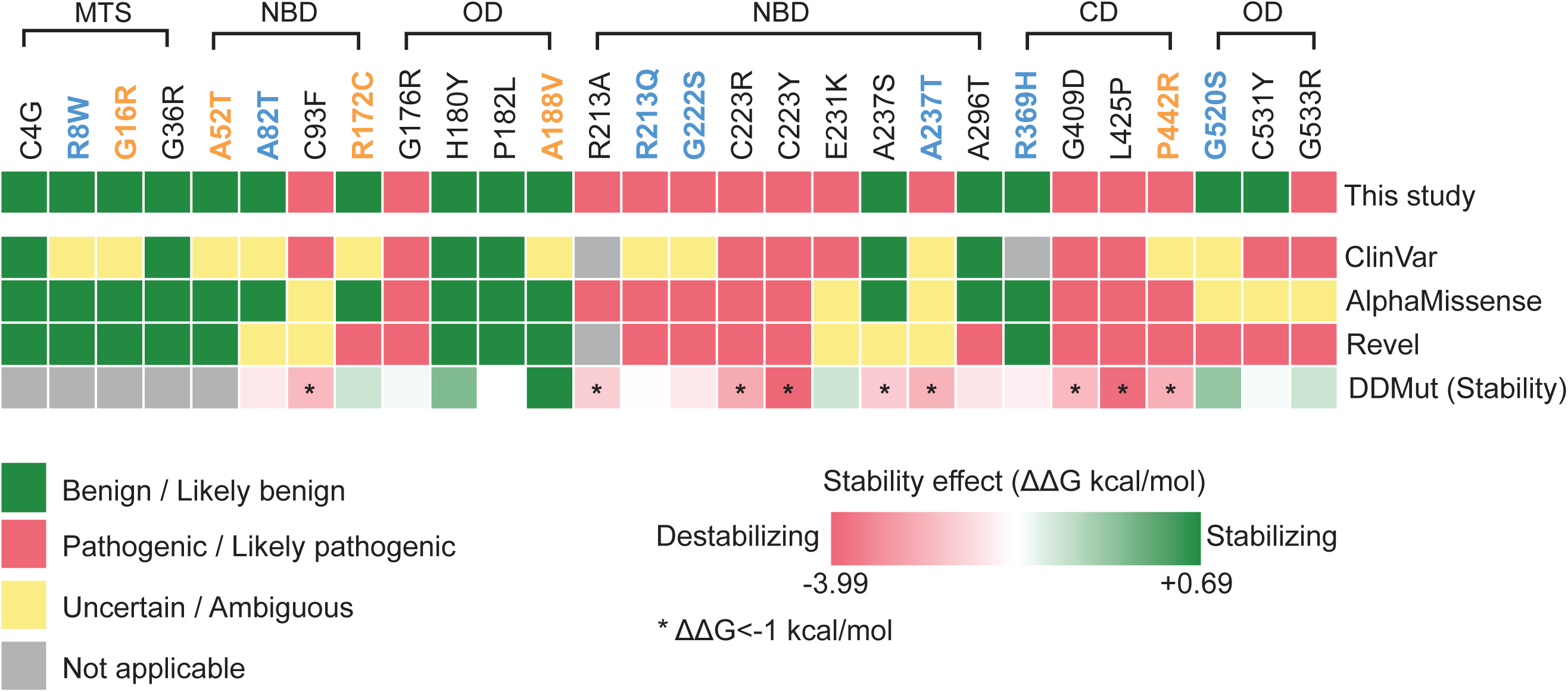
Variant classification and comparison with predictive tools. Each square represents the prediction or annotation (ClinVar) of the respective sources. Cut-off values for REVEL are benign <0.4, pathogenic >0.8 (Wilcox et al., 2022). For AlphaMissense, benign <0.34, pathogenic >0.564 (Tordai et al., 2024). For DDMut, the change in Gibbs Free Energy (ΔΔG) is indicated as stabilizing (positive) or destabilizing (negative). Values less than -1 are considered strongly destabilizing (asterisks). Blue variant labels indicate ASD, orange labels indicate VUS.

We compared our classifications to the computational predictors AlphaMissense and REVEL (Figure 4) (Cheng et al., 2023; Ioannidis et al., 2016). AlphaMissense predictions did not contradict any of our classifications, however we were able to provide benign/pathogenic classifications for six variants that were predicted to be uncertain by AlphaMissense. AlphaMissense predictions also did not contradict any ClinVar benign or pathogenic classifications. REVEL predictions also largely agreed with our classifications as well as with ClinVar. However, REVEL predicted one ClinVar benign as pathogenic (A296T), which also contradicted our benign classification. REVEL also disagreed with three more of our classifications (R172C, G520S and C531Y), where in all three cases REVEL predicted pathogenicity whereas our assay classified as benign.

To gain additional insight into underlying mechanisms of pathogenicity we computed variant stabilities using the program DDMut, which models the variant amino acid into the SSADH structure and predicts the resulting change in ΔΔG (Figure 4) (Zhou et al., 2023). ΔΔG values less than -1.0 are generally considered to be strongly destabilizing, whereas values between -1.0 and -0.5 moderately, and between -0.5 and 0 weakly destabilizing. DDMut predicted a number of variants to be strongly destabilized, including C93F, R213A, C223R, C223Y, A237S, A237T, G409D, L425P and P442R. All of these were found to have decreased function, except A237S. Additionally, all variants classified as benign by ClinVar or our assay were predicted to be stable, with the exception of A237S. Destabilization of the ASD variant A237T and the VUS P442R suggested this may be an underlying mechanism leading to their loss of function.

## DISCUSSION

We developed an assay in yeast that we believe mimics many of the metabolic consequences of loss of SSADH function in neurons, which we used to define the functional consequences of *ALDH5A1* variants found in ASD. Cellular models of *ALDH5A1* deficiency demonstrate mitochondrial dysfunction, oxidative carbonyl stress, mTOR hyperactivation, impaired autophagy, disrupted GABA receptor trafficking and defects in dendritic and synaptic maturation (Afshar-Saber et al., 2024; Latini et al., 2007; Menduti et al., 2020; Vogel et al., 2016; Vogel et al., 2017a; Vogel et al., 2017b). Many of these phenotypes arise from the inability to catabolize succinate semialdehyde, resulting in an increase of GHB and GABA, and altered NAD+/NADH balance. In the presence of GABA, *uga2Δ* yeast are unable to catabolize succinate semialdehyde to succinate, mirroring the effects of SSADH deficiency in patients. The cellular consequences are very similar, including increased GABA, succinate semialdehyde and GHB, altered redox capacity and additional metabolic phenotypes (Bach et al., 2009; Lakhani et al., 2014; Márquez et al., 2021; Ramos et al., 1985). These consequences are likely to be the cause of the decreased growth we observe for *uga2Δ* yeast grown in the presence of GABA. Close agreement of our assay results with the clinical designations for the calibrating benign and pathogenic variants further supports the clinical relevance of the yeast assay. We also find the yeast assay to be in better clinical agreement than *in vitro* assays of variant SSADH biochemical activity, which surprisingly find substantial decreases in activity of several benign variants. These include P182L and A237S, which are the second and fourth most common missense variants present in gnomAD. This suggests cell physiological context is important in accurately modelling *ALDH5A1* disease and that measuring biochemical activity alone may overestimate variant impact on function.

Having determined the clinical validity of our assay, we used it to probe the functionality of seven ASD variants and five VUS to make predictions regarding their pathogenicity. We predict three ASD variants (R213Q, G222S, and A237T) and one VUS (P442R) to be likely pathogenic. All three ASD variants reside in the NAD+ binding domain (NBD), while P442R resides in the catalytic domain (CD), consistent with the functional importance of these domains. Indeed, the positively charged Arg-213 residue is critical for binding of SSA (Kim et al., 2009). All calibration benign and pathogenic variants in these domains align functionally with their classifications giving us high confidence in our clinical predictions for variants in these domains. AlphaMissense similarly predicts R213Q, G222S and P442R to be pathogenic, but classifies A237T as ambiguous. Additionally, we predict with high confidence the ASD variants A82T (NBD) and R369H (CD), and the VUS A52T (NBD) and R172C (NBD) as likely benign.

In the mitochondrial targeting sequence, we predict the ASD variant R8W and the VUS G16R to be likely benign. However, no pathogenic variants have been described in this region (Table S1), hence we were unable to assay calibration pathogenics, making our confidence in these predictions lower. But the absence of pathogenic variants in this region suggests variants in this region are likely to not severely disrupt function. In the oligomerization domain we predict the VUS A188V and the ASD variant G520S to be likely benign. For this domain our functional data for two out of three calibration pathogenic variants are consistent with their ClinVar pathogenic classifications (G176R and G533R), however the C531Y variant appears to be functional, contradicting its pathogenic ClinVar classification. Unpublished data (“data not shown” in Pop et al.) indicate this variant exhibits less than 1% of reference *ALDH5A1* catalytic activity (Pop et al., 2020). Thus, the yeast assay may not always be a reliable predictor of variant function in this domain and we have lower confidence in our benign predictions for A188V and G520S. AlphaMissense predicts A188V to be benign, but predicts G520S as uncertain, suggesting G520S may need further examination.

The G222S ASD variant, along with the adjacent C223R pathogenic calibration variant are unique in that their activity is less than zero. This indicates that expressing these variants causes *uga2Δ* yeast to grow slower in the presence of GABA than yeast expressing the empty vector control. This presents the possibility that these variants are exhibiting dominant negative activity. Both residues are identical in all species from humans to *E. coli*., and individuals with variants in Cys-223 have strong SSADH deficiency phenotypes (Akaboshi et al., 2003; Di Rosa et al., 2009), suggesting increased pathogenicity of these variants. These two residues are located in a turn between two regions of secondary structure that contain residues crucial for NAD^+^ binding, hence it is conceivable that aberrant NAD binding/release could be more deleterious to the cell than the absence of the protein altogether. This has yet to be described for SSADH deficiency and may have been missed given the limitations of SSADH biochemical assays in uncovering dominant negative effects. Further studies are necessary to determine if these variants are indeed true dominant negatives and what the clinical implications may be for SSADH deficiency and ASD.

Our finding that *ALDH5A1* variants found in individuals with ASD are loss of function implies *ALDH5A1* missense mutations may be a causative factor in ASD. This is consistent with the fact that 30-60% of individuals with SSADH deficiency have a formal ASD diagnosis (Tokatly Latzer et al., 1993). SSADH is a critical enzyme in GABA degradation and alterations in GABA catabolic flux lead to altered excitatory/inhibitory balance, which is hypothesized to be a central aspect of some forms of ASD (Hollestein et al., 2023; Nelson and Valakh, 2015). ASD is linked to genes regulating GABAergic signalling including ABAT, GAD1/2, SLC6A1, GABRB3, and GABRA5 (Goodspeed et al., 1993; Rodriguez-Gomez et al., 2021; Zhao et al., 2022). ABAT encodes for GABA transaminase and individuals with ABAT mutations also show ASD associated phenotypes (Barnby et al., 2005). Diminished mitochondrial redox capacity and increased mTOR signalling are also important mechanisms underlying ASD. Individuals with ASD with *ALDH5A1* mutations are likely heterozygous (we found no record of compound heterozygosity), suggesting that ASD phenotypes in these individuals may represent a milder form of SSADH deficiency, given the presence of a functional copy of *ALDH5A1*. This is consistent with there being lower levels of GHB in individuals diagnosed with SSADH deficiency with ASD. Genetic context may also be important, where individuals with ASD but not SSADH deficiency require a second mutation in genes linked to GABAergic signalling, mitochondrial redox and/or mTOR signalling for ASD to occur.

Finally, it is interesting to note that four of the seven ASD variants tested in this study were functional (R8W, A82T, R369H and G520S), even though *ALDH5A1* is considered a high confidence ASD gene by SFARI. Three out of four of the functional ASD variants are designated as VUS in ClinVar (one is not present), and AlphaMissense or REVEL make either incorrect or uncertain predictions for two of the four, thus stressing the need for clinically calibrated functional assays for elucidating the true genetic causes of ASD.

## DATA AND RESOURCE AVAILIBILITY

All relevant data and details of resources can be found within the article and its supplementary information.

## AUTHOR CONTRIBUTIONS STATEMENT

Conceptualization: BY, SR, JC, PP, DA, CL. Methodology: BY, SR, JC, PP, DA, CL. Software: SR, PP. Validation: BY, CL. Formal analysis: BY. Investigation: BY, IR, JS, BW, JL, SR. Resources: BY, SR, PP, CL. Data Curation: BY, SR. Writing – original draft preparation: BY, CL. Writing – review and editing: BY, SR, PP, DA, CL. Visualization: BY. Supervision: BY, PP, DA, CL. Project administration: JC, PP, DA, CL. Funding acquisition: JC, PP, DA, CL.

